# Ancestra: A lineage-explicit simulator for benchmarking B-cell receptor repertoire and lineage inference methods

**DOI:** 10.64898/2026.04.17.718988

**Authors:** Reza Hassanzadeh, Nika Abdollahi, Sofia Kossida, Véronique Giudicelli, Changiz Eslahchi

## Abstract

High-throughput B-cell receptor sequencing has transformed the analysis of adaptive immunity, but benchmarking clonal grouping and lineage reconstruction methods remains limited by the absence of datasets with known evolutionary histories. Here we present Ancestra, a lineage-explicit simulator of B-cell receptor heavy-chain affinity maturation. Ancestra models stochastic V(D)J recombination, context-dependent somatic hypermutation, affinity-based selection and clonal expansion while recording complete parent-child relationships and mutation events. The framework generates BCR heavy-chain sequence datasets together with their corresponding ground-truth lineage trees, enabling direct benchmarking of lineage-aware analytical methods. Across simulations, Ancestra recapitulates key properties of human repertoires, including complementarity-determining region 3 length distributions, amino-acid usage patterns, junctional mutation patterns consistent with IMGT criteria and heterogeneous branching topologies. Simulated lineages also reveal multi-label lineage trees, in which identical nucleotide sequences can arise independently along distinct evolutionary paths. Ancestra provides a practical foundation for rigorous benchmarking of lineage-aware immune repertoire analysis.

## 1. Introduction

Advances in high-throughput immune repertoire sequencing (AIRR-seq) have enabled detailed characterization of B-cell receptor (BCR) repertoires across diverse biological and clinical contexts, including vaccination, infection, autoimmunity, and cancer [1-3]. These technologies provide unprecedented access to the diversity, clonal structure, and dynamics of adaptive immune responses, allowing large-scale analysis of V(D)J recombination patterns, somatic hypermutation (SHM), and clonal expansion at nucleotide and amino-acid resolution. At the same time, AIRR-seq data have highlighted substantial methodological challenges in disentangling stochastic receptor generation, context-dependent mutation, affinity-based selection, and clonal evolution from sequence data alone.

Affinity maturation of B cells in germinal centers is an inherently evolutionary process, driven by iterative cycles of mutation and selection that generate complex, branching clonal lineages rather than linear optimization trajectories [4, 5]. During this process, low-affinity naïve BCRs evolve into high-affinity antibodies through SHM and competitive selection, producing characteristic lineage topologies and expansion patterns. Understanding these dynamics therefore requires models that explicitly capture evolutionary trajectories and lineage structure, rather than relying solely on aggregate repertoire-level statistics.

A key determinant of lineage structure and evolutionary trajectories in germinal centers is the mutational process that fuels B-cell diversification. Somatic hypermutation introduces point mutations into immunoglobulin coding sequences in a strongly context-dependent manner, with well-characterized hotspot motifs (e.g. WRC, GYW, DGYW and WRCH) exhibiting elevated mutability and cold-spot motifs (e.g. SYC, GRS) showing suppressed mutation rates. These sequence-context biases generate a highly non-uniform mutational landscape that constrains accessible evolutionary pathways and interacts with affinity-based selection and structural constraints of the antibody molecule. The interplay between context-dependent mutation, selection, and clonal competition ultimately shapes lineage expansion, tree topology, and the distribution of functional antibodies observed in mature repertoires.

The combined effects of context-dependent mutation, affinity-based selection, and stochastic clonal expansion give rise to highly complex and heterogeneous BCR repertoires, whose underlying evolutionary processes cannot be directly observed from sequence data alone. Computational simulators have thus become essential tools for studying BCR repertoire dynamics, benchmarking analytical methods, and exploring hypotheses that are difficult or impossible to address experimentally. Existing simulators such as immuneSIM and AIRRSHIP generate synthetic repertoires that reproduce key statistical properties of experimental data, including V(D)J gene usage, CDR3 length distributions, and aggregate SHM patterns [1,6]. These tools have been widely adopted for validating repertoire analysis pipelines, benchmarking clonal grouping strategies, evaluating diversity metrics, and supporting machine-learning–based approaches for immune repertoire analysis [2].

More recently, simulation frameworks have been extended to support the development and benchmarking of machine-learning methods applied to AIRR data. In this context, Chernigovskaya et al. introduced LigO [7], a flexible simulation suite designed to generate AIRR datasets with implanted immune signals for receptor- and repertoire-level machine-learning benchmarking. LIgO enables controlled signal implantation while preserving repertoire generation statistics and provides scalable synthetic datasets for evaluating classification and representation-learning methods. This work represents an important advance toward principled benchmarking of AIRR-ML approaches and highlights the growing role of simulation as a source of ground-truth data in immunogenomics.

Despite these advances, current simulation frameworks, including ML-oriented simulators, largely decouple immune signal implantation and repertoire statistics from explicit models of B-cell evolutionary dynamics. In particular, most simulators output collections of sequences without native representations of somatic hypermutation along phylogenetic lineages, affinity-dependent selection, or the clonal tree structures that emerge during germinal center reactions. Lineage relationships are typically reconstructed post hoc using clustering or phylogenetic inference methods applied to simulated or experimental sequences. Benchmarking studies have shown that inferred clonal grouping and lineage trees can vary substantially depending on algorithmic choices, distance metrics, and modeling assumptions, complicating objective assessment in the absence of known parent–child relationships [8–10].

This limitation is particularly consequential as lineage-aware analyses are increasingly used to study affinity maturation, clonal competition, and immune memory. Experimental studies emphasize the complex, branching nature of BCR evolution in germinal centers, while computational analyses demonstrate that lineage inference remains under-constrained when ground-truth evolutionary histories are unavailable [4, 5, 10]. Together, these observations reveal a methodological gap between biologically motivated evolutionary questions and the simulation tools available to evaluate lineage-aware analytical methods.

Here, we present Ancestra, a lineage-explicit computational framework for simulating B-cell receptor evolution. Ancestra models the full process from naïve repertoire generation through somatic hypermutation, affinity-based selection, and clonal expansion, while explicitly recording mutation events and parent–child relationships across generations. The framework implements generation-by-generation simulation of mutation and selection, enabling full traceability of evolutionary events along clonal lineages. Each simulation produces nucleotide and amino-acid sequences, clone abundances, affinity trajectories, and complete lineage trees with known ground truth, formatted in accordance with current AIRR data standards [11].

By validating Ancestra against established experimental repertoire statistics and by analyzing lineage-level structural properties, we demonstrate that the framework generates biologically plausible repertoires while enabling rigorous benchmarking of lineage-aware analytical methods. This lineage-explicit perspective provides a foundation for systematic evaluation of evolutionary inference approaches and for investigating how mutation and selection jointly shape BCR evolutionary trajectories.

## 2. Methods

### 2.1. Overview

Ancestra models the generation and evolution of B-cell receptor (BCR) heavy chain sequences through a multi-step computational framework, comprising: (i) stochastic V(D)J recombination to produce a single naïve BCR heavy-chain rearrangement. By default, each replicate is initialized from a single naïve ancestral heavy-chain sequence to generate a ground-truth lineage; however, the simulator supports multiple ancestral sequences (and/or multiple independent replicates) to produce sets of lineages when desired; (ii) iterative rounds of nucleotide-level somatic hypermutation and translation, (iii) amino-acid CDR3-based affinity evaluation to one or more antigen epitopes, and (iv) generation-wise selection and branching (clonal expansion) over discrete generations. Each simulation returns the set of generated nucleotide sequences, their translated amino-acid sequences, per-sequence affinity scores, parent–child relationships and sequence abundances. Unlike many simulators that only output a set of BCR sequences, Ancestra additionally produces an explicit BCR lineage tree that records parent–child relationships and branch lengths so that each sequence’s evolutionary path and the mutation/selection events that produced it are unambiguously traceable. The following subsections describe each step of the simulation framework in detail.

### 2.2. Input data and notation

Let 𝒱, 𝒟 ,𝒥 denote the user-provided sets of germline *V, D* and *J* nucleotide sequences. Let *ε* = {*E*_1_,…, *E*_m_ } denote the set of antigen epitope amino-acid sequences used for affinity evaluation. Scalar parameters that control the simulation include the per-base baseline mutation rate *µ*, the maximum number of generations G_*max*_, the minimum and maximum generation affinity thresholds T_*min*_, T_*max*_, and a small set of hotspot/coldspot motif weights.

### 2.3. Naïve BCR heavy-chain generation

A single naïve BCR heavy chain is produced by the following procedure:

1. Segment sampling: One *V*, one *D* and one *J* segment are sampled uniformly at random from the sets 𝒱, 𝒟, and 𝒥, respectively. By default, *V, D*, and *J* segments are sampled uniformly; however,Ancestra supports user-defined, non-uniform sampling distributions to accommodate empirical gene usage biases when available.
2. Junctional diversity: Nucleotide trimming is applied to the 3^′^ end of the *V* segment, both ends of the *D* segment, and the 5^′^ end of the *J* segment. Trim lengths are drawn from discrete probability distributions dependent on local A/T content and biased toward short deletions [12]. For a segment tail of length up to L, the probability of trimming *k* bases is given by:

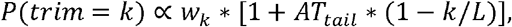

where *w*_0_ = 0.6, *w*_*k*_ = 0.2 for 1 ≤ k ≤ 4, and w_k_ = 0.01 for larger k. The local A/T fraction AT_*tail*_ ∈[0,1] increases the weight of shorter trims near A/T-rich tails. Trim lengths are sampled independently for each segment end. This formulation biases toward zero or small trims while allowing longer deletions with low probability.
3. Non-templated nucleotide insertion (N-regions): Non-templated nucleotides are added between the *V*-*D* and *D*-*J* junctions. The *N*-regions between *V* and *D* is denoted *N*_1_, and between *D* and *J* as *N*_2_. Each N-regions length is sampled uniformly from {0, … ,10}, and each inserted nucleotide is drawn uniformly from {A, C, G, T}. This formulation does not explicitly distinguish P-region (palindromic) nucleotides generated during hairpin resolution; instead, all junctional additions are treated as non-templated insertions. The insertion-length and nucleotide-frequency distributions are simplified for computational efficiency and may be replaced with empirically derived alternatives without affecting downstream mutation, selection, or lineage reconstruction procedures.
4. Sequence assembly and translation: The trimmed segments and *N*-regions are concatenated in the order *V*-*N*_1_ -*D*-*N*_2_ - *J* to form the full coding nucleotide sequence s. While minor variations in recombination architecture exist in nature, this sequence represents the standard assembly for functional BCR heavy chains. Subsequently, s is translated in all three forward reading frames; the frame containing the minimum number of internal stop codons is selected to maximize the generation of productive rearrangements. This heuristic does not model the biological frame-selection process itself, but serves to restrict the simulated repertoire to functional BCR sequences prior to somatic hypermutation. If no reading frame yields an acceptable structure (e.g. internal stops that violate acceptance criteria), the draw is rejected and the process repeats. Canonical CDR3 regions are enforced by requiring a conserved cysteine (C) at the end of the V gene segment and a tryptophan (W) residue at the start of the J segment consistent with heavy-chain structural constraints. Candidates failing these structural constraints are rejected.

### 2.4. Somatic hypermutation

Mutations at the nucleotide level are modeled probabilistically in each sequence. Regions annotated as hotspots are assigned a higher mutation probability than baseline regions, whereas coldspots are assigned a lower probability. To capture the known biological bias of mutation types, the model favors transition mutations over transversions (for example, with a 70% to 30% probability split, respectively). The implementation of this step is described in detail below.

#### 2.4.1. Hotspot and coldspot context scoring

Experimental studies consistently demonstrate that mutation rates within SHM hotspot motifs (such as WRC, GYW, DGYW, and WRCH) are substantially elevated relative to non-hotspot regions [13–17]. Empirical measurements indicate differential mutation propensities among these motifs, with DGYW/WRCH contexts generally exhibiting stronger activity than WRC-type motifs.

In Ancestra, SHM is modeled using a curated library of nucleotide-level hotspot motifs representing sequence contexts with elevated mutation probability. Each motif is assigned a weight reflecting its relative contribution to local mutation propensity. Within each motif, a designated internal target specifies the nucleotide preferentially deaminated by activation-induced cytidine deaminase (AID) [13–17]. For example, in WRC motifs, the cytosine serves as the primary AID target.

For each nucleotide position, the algorithm scans the surrounding sequence to identify motif instances whose internal target offsets align with the position of interest. The weights of all matching motifs are summed to generate a positional hotspot score that quantifies local mutation likelihood.

A complementary set of coldspot motifs (e.g., SYC and GRS) is incorporated to model reduced mutability. Coldspots impose multiplicative penalties on mutation probability at the target nucleotide and within a defined flanking window, thereby representing localized suppression of SHM. Motif matching supports IUPAC nucleotide codes, enabling detection of degenerate sequence patterns.

All SHM calculations in Ancestra are performed at the nucleotide level prior to translation. Context-dependent mutation probabilities are computed independently of amino acid translation or affinity evaluation. Default hotspot weights and coldspot penalties are provided (Table 1, Section 3.1), while users may modify existing motifs or introduce custom patterns to tailor SHM modeling. This context-based scoring framework provides a probabilistic abstraction of SHM targeting rather than a mechanistic simulation of AID biochemistry.

**Table 1.** Baseline simulation parameters.

|  | Parameter Name | Value | Description |
| --- | --- | --- | --- |
| 1 | $T_{max}$ | 0.8 | Upper bound for selecting high-affinity BCRs during simulation |
| 2 | $T_{min}$ | 0.3 | Minimum affinity required for BCRs to survive early generations |
| 3 | $G_{max}$ | 10, 15, 20, 25, 30 | Maximum depth of lineage expansion |
| 4 | $\mu$ | 0.001 | Probability of baseline mutation per nucleotide per sequence |
| 5 | n_runs | 10 | Number of independent simulation replicates |
| 6 | min_uniq_seq | 15 | Minimum number of unique nucleotide sequences required to validate a run |
| 7 | Hotspots weights | WRC: 4, GYW: 8, DGYW: 10, WRCH: 10 | Relative mutation likelihood at hotspot motifs during somatic hypermutation |
| 8 | Coldspots penalties | SYC: 0.6, GRS: 0.5 | Penalty factors reducing mutation probability at coldspot motifs |
| 9 | $p_{trans}$ | 0.7 | Bias in nucleotide substitution favoring transitions over transversions |
| 10 | $p_{sub}$ | 0.3 | Survival probability for in-frame low-affinity sequences |
| 11 | $p_{stop}$ | 0.005 | Base survival probability for high-affinity sequences with stop codons |
| 12 | $p_{min}$ | 0.001 | Base survival probability for low-affinity sequences with stop codons |
| 13 | use_full_BCR | False | Affinity calculations should use the full BCR sequence (T) or only the CDR3 region (F). |

#### 2.4.2. Per-site mutation probability

The per-site mutation propensity is then obtained by combining the baseline per-base mutation rate *µ* with the context-dependent modifiers. If no hotspot motifs contribute to a position its mutation probability equals the baseline. If hotspots are present, the baseline is multiplied by the hotspot score. Any coldspot penalties identified within the relevant window are then applied multiplicatively to reduce the resulting probability. Finally, a single uniform random draw determines whether a mutation occurs at the target nucleotide. The nucleotide is mutated if the draw falls below the context-adjusted probability. When a nucleotide substitution is accepted at a target position, the replacement nucleotide is sampled according to a transition bias, where transitions occur with probability p_*trans*_ and transversions with probability 1–p_*trans*_. For all simulations presented here,we use p_*trans*_ = 0.7, though this parameter can be configured by the user.

### 2.5. Affinity evaluation

Affinity between a BCR and each epitope E_*k*_ in ε is computed at the amino-acid level using local pairwise sequence alignment. Let A denote the CDR3 amino-acid subsequence of the BCR. For each epitope *E*_*k*_, a local alignment score s(A, *E*_*k*_) is calculated using a substitution matrix equivalent to BLOSUM62. The raw alignment score is subsequently normalized to the interval [0, 1] to provide a comparable relative affinity measure across sequences of varying lengths. The affinity of a given BCR is defined as the maximum normalized alignment score across all epitopes in *ε*.

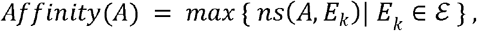

where *ns*(*A, E*_k_) denotes the normalized alignment score of *s*(*A, E*_*k*_). To improve computational efficiency during large-scale simulations, alignment scores for identical sequence pairs are cached, thereby avoiding redundant recalculations across generations.

The use of a CDR3-focused, alignment-based affinity proxy in Ancestra is a deliberate modeling choice motivated by the following considerations:

1. **Primary objective of the simulator:** Ancestra is designed to model B-cell evolution rather than provide precise biophysical binding predictions. The goal is to generate realistic evolutionary trajectories with explicit parent–child relationships and known ground-truth lineage trees.
2. **Computational tractability:** Each simulation evaluates millions to billions of candidate sequences. Using complex structural modeling or physics-based predictions at this scale would make large-scale simulations impossible due to the extreme computational cost.
3. **Requirement for relative ranking:** Affinity maturation depends on the relative competition between clones. A stable and consistent scoring proxy is sufficient to drive these selection and branching dynamics without needing absolute binding energy values.
4. **Biological relevance of the CDR3:** The CDR3 region contains the most significant functional diversity for antigen recognition. Focusing on this region captures the necessary signals for competitive selection while maintaining efficiency.
5. **Empirical plausibility:** Even with this approximation, the simulated repertoires successfully reproduce the statistical properties found in real BCR datasets. This confirms that the proxy effectively generates biologically plausible evolutionary patterns.

In light of these considerations, the affinity module in Ancestra is fully modular and replaceable. Users may compute affinity using either CDR3-only or full BCR sequences, and alternative scoring models can be incorporated without modifying lineage tracking or tree construction. This design ensures that more detailed biophysical models can be adopted when computational resources permit, while preserving the scalability required for large-scale lineage-explicit simulations.

### 2.6. Selection and clonal expansion

Evolution proceeds in discrete generations *g*= 0,1, …, G_*max*_. By default, generation *g* = 0 contains a single naïve BCR heavy-chain sequence generated in the initialization step described above; however, the framework supports initialization from multiple naïve founders or repeated independent replicates to generate collections of lineages when desired. All selection and survival probabilities are parameterized and user-configurable, allowing the strength and stringency of selection to be systematically varied across simulations. The per-generation dynamics are defined as follows:

#### 2.6.1. Descendant production and selection

In each generation, every surviving B cell independently attempts to produce two descendants. For each attempted descendant, a candidate nucleotide sequence is generated by the mutation model, translated, and inspected for the presence of the canonical CDR3 region. Candidates that lose the CDR3 region are discarded as non-functional. For retained candidates we record the number of internal stop codons and evaluate CDR3 affinity against the antigen set.

#### 2.6.2. Affinity thresholds and selection

A generation-dependent affinity threshold is defined that increases gradually from an initial value T_*min*_ to a final value T_*max*_ across generations according to a user-specified schedule. This schedule controls the progressive stringency of selection as evolution proceeds and is fully parameterized. For each candidate descendant, survival is determined probabilistically based on its affinity relative to the current generation threshold and on sequence functionality:

- If the candidate’s affinity meets or exceeds the generation threshold and the translation is in-frame (i.e., free of internal stop codons), survival is deterministic.
- If the candidate is in-frame but its affinity falls below the threshold, it is retained with a moderate probability p_*sub*_, a user-defined parameter controlling permissive survival below the threshold.
- Candidates containing internal stop codons are strongly penalized. High-affinity candidates with stop codons survive with a small probability 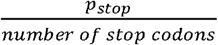, which decreases with the number of stop codons, whereas low-affinity candidates with stop codons have a negligible survival probability 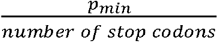. Both p_*stop*_ and p_*min*_ are user-defined parameters that control the stringency of selection against premature termination.

For each candidate, a single independent random draw determines survival by comparing the draw to its assigned probability. Surviving descendants enter the next generation with an initial abundance of one, and their parent identifiers are recorded to enable lineage reconstruction. All probabilities and threshold parameters are user-configurable and can be systematically varied to explore different selection regimes.

#### 2.6.3. Termination

The iterative process continues until either the predefined maximum number of generations (G_*max*_) is reached or no descendants survive into the next generation (population extinction). Termination criteria are parameterized, enabling flexible control over lineage depth and repertoire size. Optionally, independent replicates can be generated until user-defined acceptance criteria are met, such as achieving a minimum number of unique sequences in the final repertoire or obtaining at least one sequence meeting the maximal affinity threshold T_*max*_.

### 2.7. Post-processing: abundance consolidation and tree construction

After completion of the branching simulation, a post-processing step is applied to consolidate abundances and construct a lineage tree representation while preserving all mutation and parent– child information generated during simulation.

First, any node that is identical in nucleotide sequence to its parent (i.e., with zero recorded substitutions) is removed, and its abundance is transferred to the parent. Child links of the removed node are reconnected to the parent. This step eliminates trivial duplication events arising from accepted non-mutational offspring and does not alter lineage topology or mutation history. Additionally, sequences with identical nucleotide sequences that descend from the same parent are collapsed into a single representative entry, with its abundance set to the sum of the merged sequences’ abundances. Parent identifiers are then reassigned to preserve the integrity of the lineage tree structure.

For lineage tree construction, branch lengths between a child and its parent are set equal to the Hamming distance (i.e., the number of nucleotide substitutions) between the two sequences. This representation encodes the realized mutational distance along each edge and is intended for lineage-aware benchmarking rather than phylogenetic inference under an explicit evolutionary model. The tree root corresponds to the naïve BCR sequence. Alternative branch-length definitions can be substituted without affecting the recorded lineage structure.

### 2.8. Output

For each accepted simulation replicate, Ancestra produces three primary outputs:

i. Clone summary table: A tabular file listing all generated sequences with associated metadata, nucleotide and amino-acid sequences, CDR3 coordinates, generation index, parent identifier, mutation count, affinity score, and abundance.
ii. FASTA sequence file: A FASTA-formatted file containing all nucleotide sequences, with sequence abundances encoded in the header lines.
iii. Lineage tree file: A text file in Newick format representing the generated BCR lineage tree, with branch lengths corresponding to the number of nucleotide substitutions between parent and child nodes and the root corresponding to the naïve BCR sequence.

All outputs preserve explicit parent–child relationships and are designed to facilitate downstream lineage-aware analysis, benchmarking, and visualization using standard AIRR-compatible tools.

## 3. Results

In this section, we present a systematic evaluation of the repertoires generated by Ancestra. Our primary motivation for developing this tool was to generate benchmark datasets for evaluating methods that reconstruct lineage trees from BCR sequence collections. By “benchmark dataset,” we refer to a dataset that includes both the BCR sequences and their corresponding ground-truth lineage trees, enabling direct assessment of reconstruction accuracy. The absence of such paired data had long been a major limitation, making it difficult to compare existing reconstruction methods or assess their accuracy in a controlled setting. Using Ancestra, as described in the Methods section, we are able to generate benchmark datasets. However, our contribution extends beyond data generation alone. We also assess whether the simulated BCR sequences reproduce key statistical, structural, and mutational properties observed in empirical human B-cell receptor datasets. Accordingly, we conducted a comprehensive set of quantitative analyses on the simulated repertoires. Specifically, we examined the length distribution of CDR3 regions, amino acid frequency compositions, patterns of somatic hypermutation, and compliance of junctional mutation patterns with IMGT-defined constraints across simulated lineages.

### 3.1. Input Data and Parameter Configuration

Unless otherwise stated, all results presented in this section were generated using a common set of simulation parameters, summarized in Table 1. These parameters were selected to balance biological plausibility, computational efficiency, and sufficient evolutionary depth for lineage-aware analyses. While Ancestra allows full user control over all parameters, fixing a baseline configuration enables consistent comparison across simulations and figures.

We varied the maximum number of generations (*G*_*max*_ ∈ {10,15,20,25,30}) to examine lineage properties across different evolutionary depths. For each *G*_*max*_ value, 10 independent simulation replicates were performed. All other parameters, including mutation rate, affinity thresholds, and motif-specific mutation modifiers, were held constant across runs.

Input germline gene sets (𝒱, 𝒟 and 𝒥) were obtained from IMGT/GENE-DB and restricted to functional human IGH genes [18], and antigen epitopes set E were drawn from the Immune Epitope Database (IEDB) with a focus on influenza-related sequences [19]. This standardized parameterization ensures that differences observed across conditions reflect changes in evolutionary depth rather than shifts in underlying simulation settings.

### 3.2. Length distribution of CDR3s

Previous studies have reported that human CDR3 lengths typically fall between 8 and 25 amino acids, with rare cases extending to shorter or longer extremes (e.g., lengths as short as 4 or as long as 34) [20]. Using the baseline parameter settings described above, we examined the distribution of CDR3 lengths across all simulated sequences (Fig. 1). As illustrated in the leftmost violin plot of Figure 1, the aggregated distribution aligns closely with the expected human range, exhibiting a median CDR3 length of approximately 18–20 amino acids. We then conducted the calculations and analyses separately for each *G*_*max*_. Figure 1 demonstrates that the resulting distributions remain centered within the biologically relevant range, with median values consistently comparable to those of the overall dataset. Rare outliers with unusually short or long CDR3 regions were observed at multiple generation depths (e.g., *G*_*max*_ =15 and *G*_*max*_=25), consistent with the low-frequency extremes reported in empirical human repertoires. Together, these results indicate that Ancestra preserves realistic CDR3 length distributions across a range of evolutionary depths.

**Figure 1.**
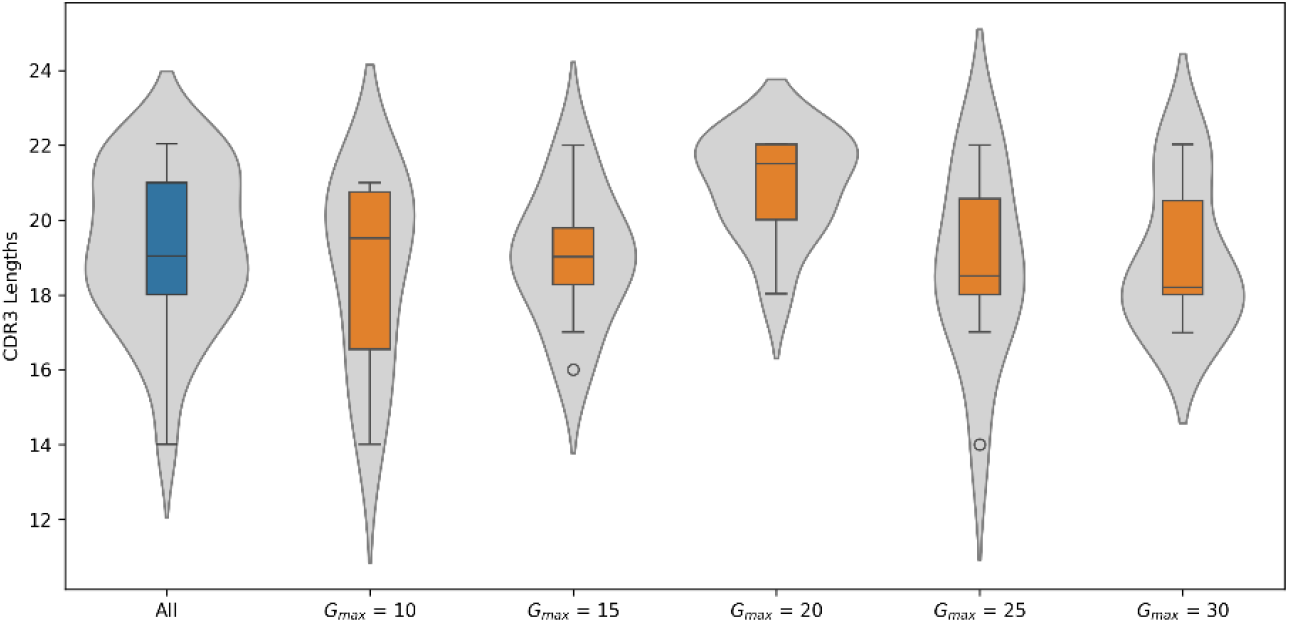
Distribution of CDR3 lengths. Violin plots showing the distribution of CDR3 amino⍰acid lengths for all simulated BCR sequences (“All”) and for each. Each violin contains an embedded boxplot indicating the median and interquartile range. All distributions fall within the typical human CDR3 length range (8–25 amino acids).

### 3.3. Amino Acid Frequency Distributions

#### 3.3.1. Simulated Sequences

To assess how amino-acid composition evolves in the simulated BCR heavy chain sequences, we examined residue frequency distributions across all simulated sequences (Figure 2) as well as within each condition (Supplementary Figure 1). Boxplots were generated separately for whole sequences and for the CDR3 region.

**Figure 2.**
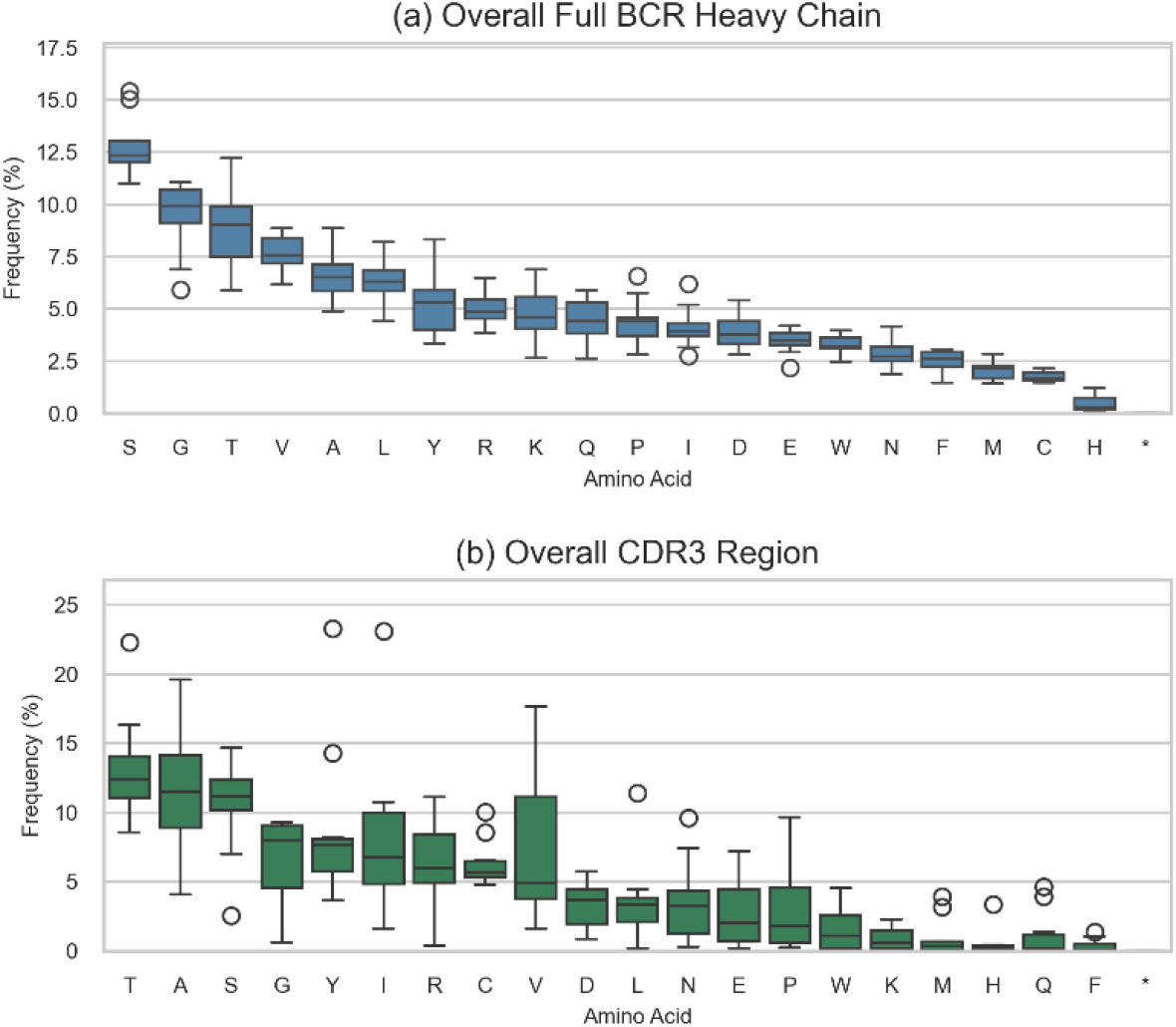
Amino Acid Frequency Distributions. **(a)** Distribution of amino-acid frequencies across full simulated BCR heavy-chain sequences. Serine (S), Glycine (G), and Threonine (T) are the most abundant residues, whereas Methionine (M), Cysteine (C), and Histidine (H) occur at minimal levels. **(b)** Amino-acid frequency distribution restricted to the CDR3 region. The CDR3 exhibits a distinct compositional profile, with Threonine (T), Alanine (A), Serine (S), and Glycine (G) dominating the region, consistent with the elevated structural flexibility and biochemical diversity characteristic of CDR3 loops. Low-frequency residues include Histidine (H), Glutamine (Q), and Phenylalanine (F). Together, these distributions illustrate region-specific compositional biases encoded by the simulation framework.

In both analyses, the overall amino-acid composition of full-length sequences remained stable across generations, exhibiting a consistent pattern dominated by residues commonly enriched in immunoglobulin variable domains.

- **Dominant Residues:** Serine (S), Glycine (G), Threonine (T), Leucine (L), Alanine (A), and Valine (V) were consistently among the most frequent residues, each typically appearing between approximately 8% and 15% of the total sequence composition.
- **Low-frequency residues and structural constraints**: Less common residues, such as Phenylalanine (F), Methionine (M), Cysteine (C), and Histidine (H), consistently appeared at very low frequencies (generally <2.5%). This pattern accurately reflects their expected limited usage in natural BCR sequences and demonstrates the simulator’s preservation of key structural constraints, including restricted incorporation of free cysteines.
- **Repertoire Stability:** The narrow interquartile ranges (IQRs) observed for most residues in the full-sequence distributions indicate that the Ancestra produces compositionally stable repertoires with limited generation-to-generation variability, suggesting rapid convergence to a biologically plausible global amino acid composition.

In contrast to full-length sequences, the CDR3 region displayed greater heterogeneity and a broader dynamic range of amino acid frequencies, which aligns with its central role in antigen recognition and functional diversification.

- **Key Residue Enrichment:** Across all conditions, residues such as Threonine (T), Serine (S), Alanine (A), Glycine (G), and Tyrosine (Y) typically showed the highest median frequencies. Enrichment of Glycine and Tyrosine is particularly notable, reflecting Glycine’s role in conferring conformational flexibility and Tyrosine’s frequent involvement in antigen-binding interfaces [9,21,22].
- **Variability and Outliers:** The CDR3 distributions displayed broader spreads and more frequent high-value outliers than the corresponding full-sequence distributions. Outliers observed for residues such as Tyrosine, Glycine, and Cysteine reflect substantial sequence-level diversity arising from junctional variation and somatic hypermutation within the hypervariable region.
- **Generational Convergence:** Although the CDR3 region exhibited higher variability overall, the shape of the amino-acid frequency distributions remained consistent across all generation depths. For example, while Glycine frequencies spanned a broad range, median values remained relatively stable (on the order of 20–30%, depending on generation depth and sampling variability), indicating that the simulation preserves core biochemical composition while allowing realistic variation within hypervariable regions.

Overall, the observed residue distribution patterns across generations qualitatively mirror those observed in empirical BCR datasets, characterized by globally stable sequence composition alongside highly variable, compositionally biased CDR3 regions. This stability supports the biological plausibility of the generated sequences and provides a consistent biochemical backdrop for subsequent lineage-level evolutionary analyses.

#### 3.3.2. Simulated Sequences vs Real Sequences

To confirm Ancestra’s ability to generate realistic B-cell receptor (BCR) sequences, we benchmarked the simulated repertoires against empirical human BCR heavy-chain data. Human BCR heavy-chain sequences were retrieved from the AntiBody Sequence Database (ABSD) and used as an empirical reference dataset (hereafter referred to as the “Real Benchmark”) [23]. Amino-acid frequency distributions in the simulated sequences were then compared to those observed in the real dataset.

As in Section 3.3.1, we first compared residue frequency distributions across all simulated sequences and the real benchmark (Figure 3), and subsequently examined these distributions separately for each condition (Supplementary Figure 2). The fidelity of the simulated sequences is illustrated by the close agreement between simulated amino-acid frequency distributions and the empirical benchmark. For the vast majority of amino acids, the benchmark frequencies (shown as red diamond markers) fall within the central density of the simulated distributions, often overlapping the interquartile range of the violin plots. This graphical agreement indicates that the simulated repertoires closely recapitulate empirical amino-acid usage patterns. To quantitatively assess similarity between simulated and real repertoires, we computed three standard divergence metrics between the empirical benchmark distribution and the median simulated distributions: Jensen-Shannon (JS) divergence, Hellinger distance, and Total Variation Distance (TVD).

**Figure 3.**
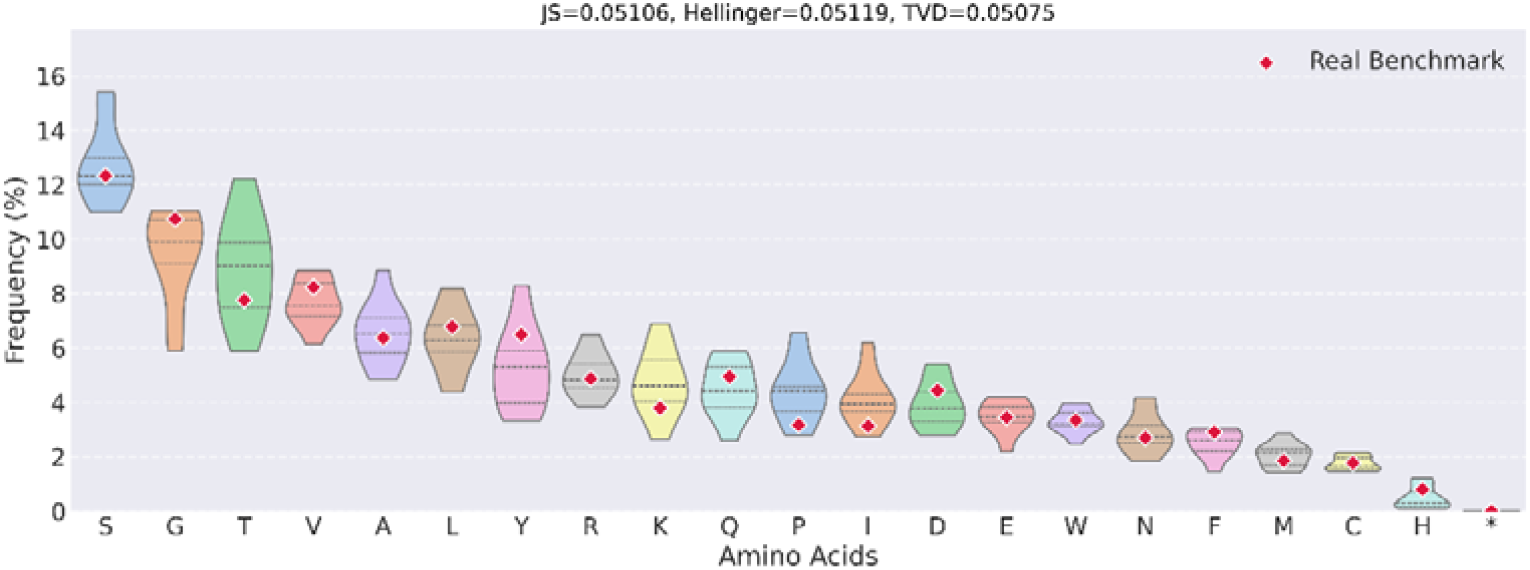
Amino-acid frequency distributions across simulated versus real BCR repertoires. Each violin represents the across-run variability of a single amino acid in the simulated repertoires, ordered from highest to lowest median simulated frequency. Red diamond markers indicate the corresponding amino-acid frequencies observed in the real benchmark dataset. Similarity between simulated and real repertoires is quantified using Jensen-Shannon divergence (JS), Hellinger distance, and Total Variation Distance (TVD), computed between the median simulated distribution and the empirical benchmark. Lower values indicate stronger agreement between simulated and observed amino-acid usage patterns.

Panel titles in Figure 3 and Supplementary Figure 2 report the divergence metrics calculated between the median simulated amino-acid frequencies and the real benchmark distribution. Quantitatively, divergence between simulated and empirical repertoires remained low across the examined evolutionary window. For the aggregate comparison shown in Figure 3, divergence values were JS = 0.05106, Hellinger = 0.05119, and TVD = 0.05075, indicating strong overall agreement.

Supplementary Figure 2 further shows that JS divergence values ranged from 0.05712 to 0.06549, Hellinger distances from 0.05614 to 0.06584, and TVD values from 0.06162 to 0.07054 across generations 10–30. Divergence decreased modestly from generation 10 to generation 25, reaching a minimum at generation 25 (JS = 0.05066; Hellinger = 0.05614; TVD = 0.06198), and remained relatively stable by generation 30. These trends suggest progressive convergence of amino-acid usage toward empirically observed distributions as simulated repertoires undergo additional rounds of mutation and selection.

Overall, these results demonstrate that Ancestra robustly reproduces amino-acid frequency patterns observed in real human BCR repertoires, with accuracy improving gradually across generations. The consistently low divergence values and stable across-run variance indicate that the evolutionary model captures key biochemical and selective constraints shaping immunoglobulin amino-acid composition. The close alignment with empirical data underscores Ancestra’s utility as a platform for benchmarking repertoire-analysis pipelines, evaluating inference methods that account for lineage and evolutionary dynamics, and generating realistic synthetic datasets to support methodological development.

### 3.4. Lineage Structure and Tree Topology

To assess whether Ancestra generates biologically plausible evolutionary structures beyond sequence-level statistics, we examined the topology and branching properties of the simulated BCR lineage trees. Representative lineage trees generated at increasing maximum generation depths (*G*_*max*_ = 10,20,30) illustrate the non-linear and heterogeneous branching structure of simulated B-cell evolution (Figure 4). Because affinity maturation proceeds through iterative rounds of mutation and selection, realistic simulation of such branching lineages is essential for lineage-aware benchmarking.

**Figure 4.**
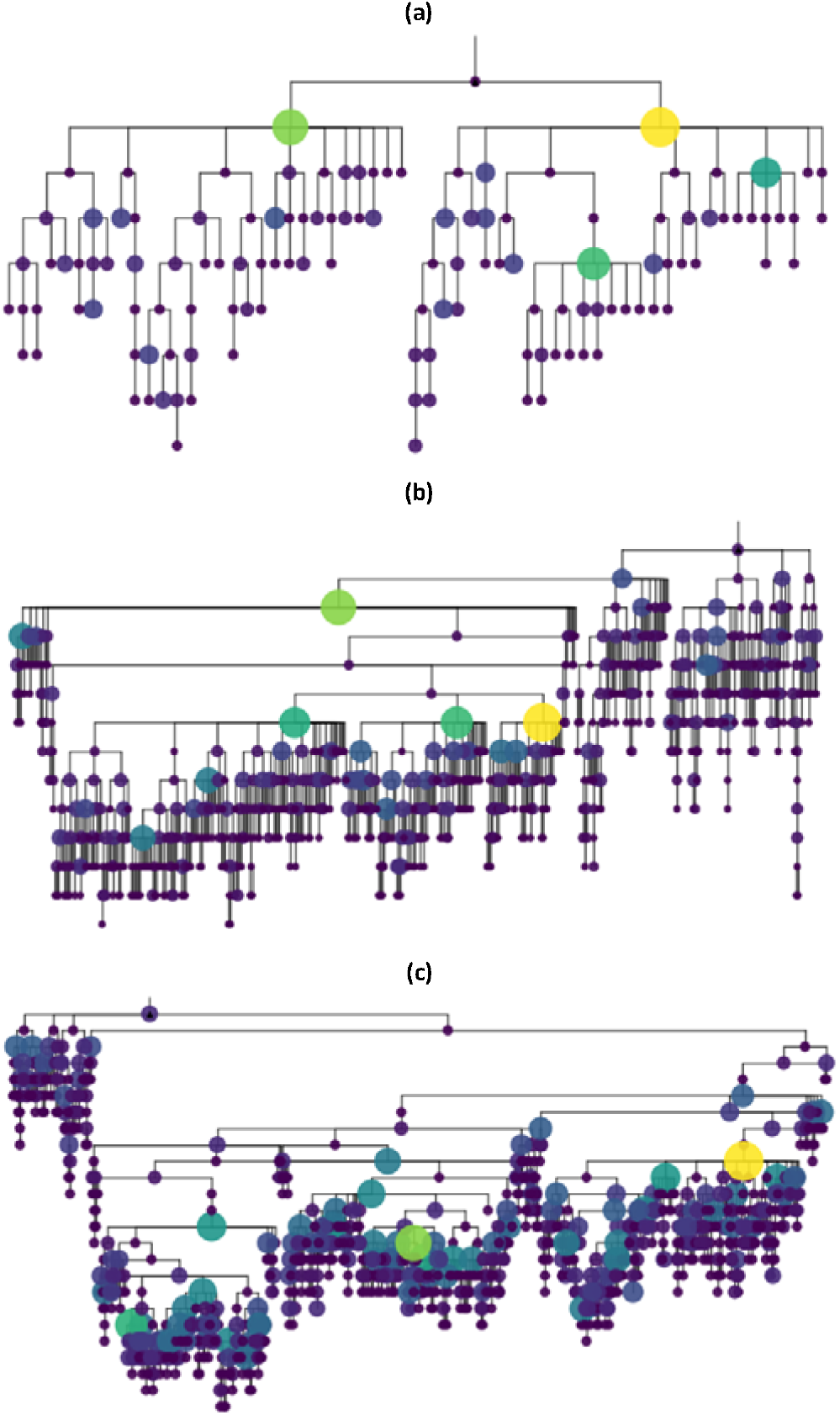
Lineage Structure and Tree Topology. Each panel shows a complete lineage tree from a single simulation replicate at different maximum generation depths: (a) ,(b) ,(c). Across increasing evolutionary depths, lineage trees exhibit heterogeneous branching and coexistence of multiple clonal branches, illustrating the non-linear structure of simulated B-cell evolution.

Across all simulation conditions, the generated lineages exhibited pronounced branching patterns, with multiple coexisting descendant clones emerging at each generation. Lineage trees were characterized by heterogeneous branch lengths and non-uniform expansion dynamics, reflecting the stochastic nature of somatic hypermutation and affinity-dependent selection. Importantly, simulated trees did not collapse into predominantly linear chains, but instead formed richly branching structures consistent with experimental observations of germinal center B-cell evolution, in which intraclonal diversification and competitive selection give rise to highly branching lineage trees rather than linear evolutionary paths [4,5].

To quantify tree structure, we examined basic topological features including total tree depth, number of branching events per generation, and the distribution of node degrees. As expected, increasing the maximum generation depth *G*_*max*_ led to deeper trees with larger numbers of terminal nodes, while preserving substantial internal branching at intermediate generations. This behavior indicates that evolutionary depth increases without inducing artificial linearization of the lineage structure.

Furthermore, the presence of competing clonal branches persisted throughout the simulation process. High-affinity clones expanded preferentially but did not completely eliminate lower-affinity branches, resulting in coexistence of multiple lineages over extended evolutionary windows. This pattern mirrors experimental evidence that germinal centers maintain diverse clonal populations rather than converging to a single dominant lineage.

Together, these results demonstrate that Ancestra produces lineage trees with realistic branching topologies and evolutionary heterogeneity, capturing key qualitative features of B-cell affinity maturation. The explicit recording of parent–child relationships and mutation events enables direct access to ground-truth lineage histories, providing a foundation for systematic evaluation of lineage reconstruction, phylogenetic inference, and evolutionary analysis methods applied to AIRR-seq data.

### 3.5. IMGT-Compatible Junction Constraints in Simulated Lineages

To validate mutation modeling in junction regions, we evaluated simulated sequences against the mutation acceptance criteria defined by IMGT/JunctionAnalysis for the 3’V-REGION, D-REGION, and 5’J-REGION. According to IMGT guidelines, unmutated (naïve) IGH sequences are required to contain zero mutations in the 3’V-REGION and 5’J-REGION and no more than two mutations in the D-REGION. For mutated (non-naïve) sequences, the corresponding accepted limits are two, four, and two mutations, respectively.

We analyzed a total of 163,585 simulated sequences, including 50 naïve sequences and 163,535 mutated sequences, generated across 50 independent simulation runs (10 replicates under five generation-depth settings). All sequences were annotated using IMGT/HighV-QUEST, and mutation counts within each junction region were extracted and evaluated against the corresponding acceptance thresholds.

All simulated sequences complied with the IMGT criteria. Among naïve sequences, 100% satisfied the acceptance limits, with mean mutation counts of 0.020, 0.160, and 0.000 in the 3’V-REGION, D-REGION, and 5’J-REGION, respectively. These small non-zero values arise from junction boundary handling during IMGT annotation rather than from biologically accepted mutations. Similarly, 100% of mutated sequences fell within the accepted limits, with mean mutation counts of 0.323, 1.448, and 1.078 in the corresponding regions.

Figure 5 compares the IMGT-accepted mutation thresholds with the observed mutation counts for naïve and mutated sequences across junction regions. Together, these results demonstrate that Ancestra produces junctional mutation patterns fully consistent with IMGT validation criteria, supporting the biological plausibility of its V(D)J recombination and junction modeling.

**Figure 5.**
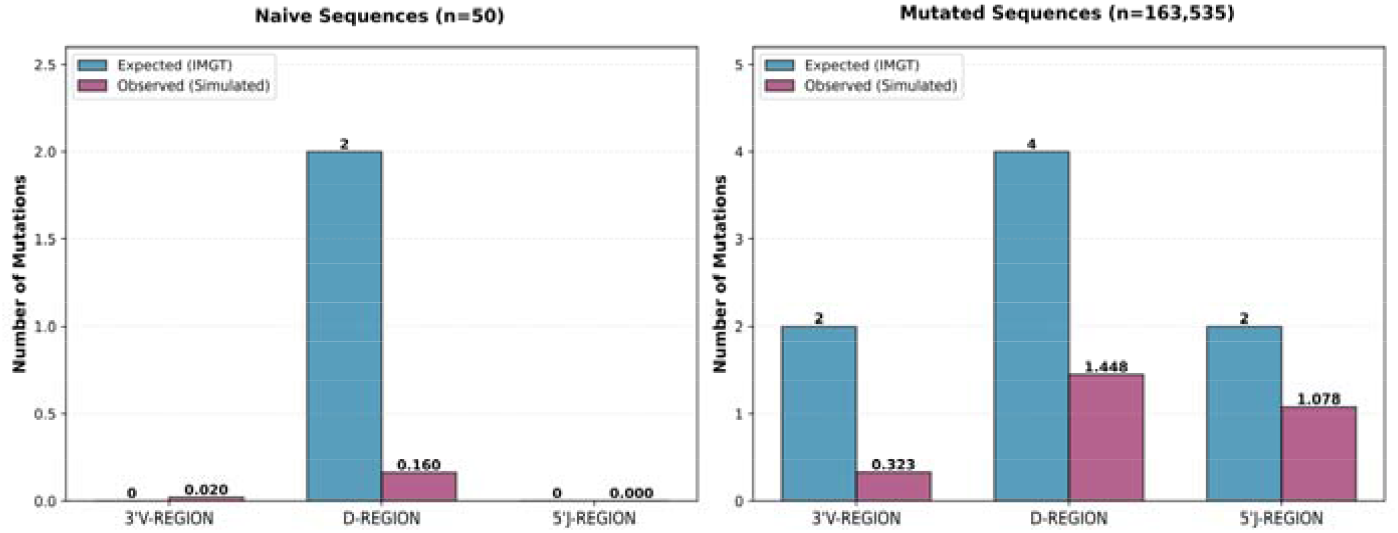
IMGT-compatible junction mutation patterns in simulated BCR sequences. Bar plots show the mean number of mutations observed in the 3’V-REGION, D-REGION, and 5’J-REGION for naïve and mutated simulated sequences. Dashed lines indicate the maximum mutation counts permitted by IMGT/JunctionAnalysis criteria for each region. All simulated sequences fall within the accepted thresholds, demonstrating full compliance with IMGT validation rules.

### 3.6. Multi⍰label Lineage Tree

During simulation of BCR heavy-chain sequences and their corresponding lineage trees, we observed cases in which identical nucleotide sequences arose independently along multiple branches of the simulated evolutionary history. Rather than mapping uniquely to a single continuous mutational trajectory, these sequence states appeared at more than one topologically distinct location within the lineage tree. We refer to this representation as a multi-label lineage tree, in which a single sequence state may correspond to multiple evolutionary paths. Representative examples of this behavior are shown in Figures 6 and 7, where nodes corresponding to identical nucleotide sequences are highlighted in red.

**Figure 6.**
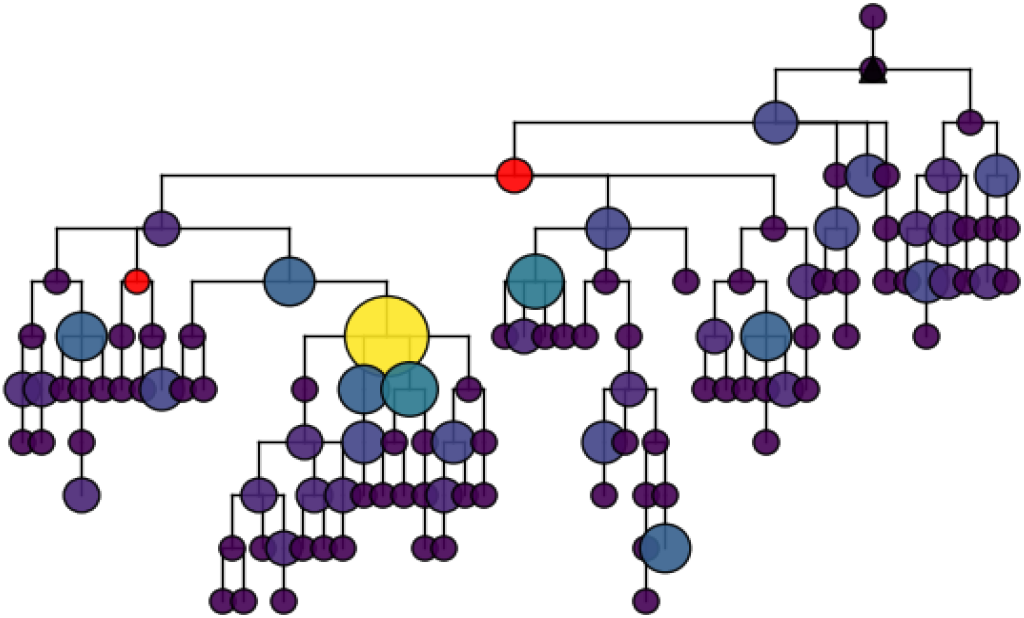
Multi-label lineage tree illustrating short-range convergent sequence states. Representative simulated BCR lineage tree exhibiting a multi-label structure, in which an identical nucleotide sequence appears at two distinct positions within the tree. Nodes corresponding to the identical sequence state are highlighted in red. In this example, the two red nodes are separated by two edges, indicating convergence over a short evolutionary distance. Branch lengths represent the number of nucleotide substitutions between parent and child nodes. This pattern reflects distinct mutational trajectories converging on the same sequence state during somatic hypermutation and selection.

**Figure 7.**
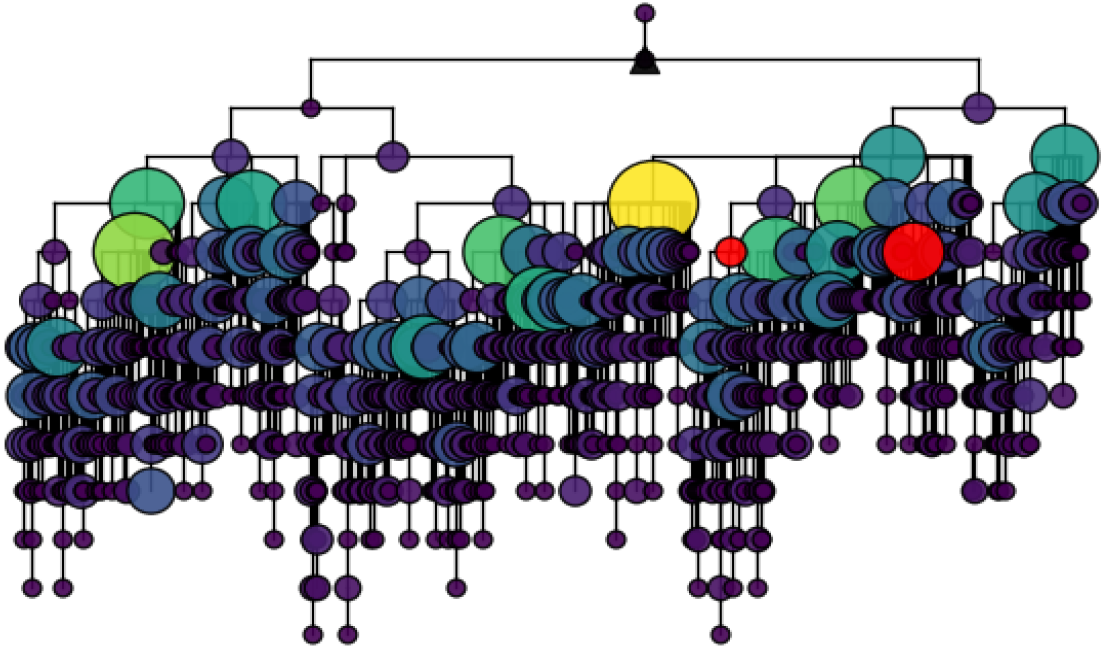
Multi-label lineage tree illustrating long-range convergent sequence states. Representative simulated BCR lineage tree showing a multi-label structure in which identical nucleotide sequences arise at topologically distant locations within the lineage. Red-highlighted nodes indicate identical sequence states separated by more than two edges, consistent with convergence occurring over more extended evolutionary paths. Branch lengths correspond to nucleotide substitution counts. This example demonstrates that identical BCR sequence states can emerge independently across distant branches of the lineage tree due to convergent evolutionary pressures.

Across all simulated datasets, multi-path sequence states were observed in approximately 40% of lineage trees. The number of such events varied across simulations: some trees contained a single instance, whereas others exhibited multiple independent occurrences. In approximately 80% of detected cases, the corresponding nodes were separated by two edges in the lineage tree (Figure 6), while in the remaining cases (e.g., Figure 7), the distance between identical sequence states exceeded two edges, indicating convergence over more distant evolutionary paths.

Conventional lineage inference frameworks typically assume that each unique sequence occupies a single node in the tree, implicitly corresponding to a single historical route of V(D)J recombination and SHM [9,10]. In contrast, the lineage-explicit simulation framework implemented in Ancestra reveals that identical sequence states can emerge through distinct mutational trajectories. This behavior reflects convergent evolution, whereby different combinations of mutation events lead to the same nucleotide or amino-acid sequence.

Such convergent sequence evolution is consistent with known biological features of B-cell affinity maturation [9,13-17,24]. Somatic hypermutation is strongly biased toward specific hotspot motifs, and selection pressures often favor particular amino-acid states that optimize antigen binding. As a result, distinct B-cell clones can independently acquire identical mutations or converge on identical sequence states. Previous studies have documented convergent patterns in BCR evolution; for example, Hoehn et al. demonstrated that unrelated B-cell lineages frequently accumulate similar or identical mutations due to shared mutational biases and functional constraints [24].

The explicit emergence of identical sequence states along multiple evolutionary paths in Ancestra highlights an additional layer of complexity in BCR lineage analysis. These results suggest that conventional single-path tree representations may obscure convergent evolutionary outcomes and underscore the value of lineage-explicit simulation frameworks with known ground-truth histories for benchmarking and method development. This phenomenon does not imply ambiguity in the simulated evolutionary history, but rather reflects distinct mutational paths converging on identical sequence states

## 4. Discussion

In this work, we introduced Ancestra, a lineage-explicit simulation framework for modeling BCR evolution that integrates stochastic V(D)J recombination, context-dependent somatic hypermutation, affinity-based selection, and clonal expansion while explicitly recording parent–child relationships and mutational histories. By preserving full evolutionary traceability, Ancestra addresses a key methodological gap in immune repertoire simulation: the absence of ground-truth lineage structures required for rigorous evaluation of lineage-aware analytical methods.

Across multiple levels of analysis, Ancestra generated repertoires that closely recapitulate known statistical and biochemical properties of human BCR sequences. Simulated CDR3 length distributions fell within empirically observed ranges and remained stable across increasing evolutionary depths, indicating that deeper lineage expansion does not distort global repertoire structure. Similarly, amino-acid composition analyses demonstrated stable full-sequence distributions alongside increased heterogeneity within the CDR3 region, reflecting the distinct functional and structural roles of framework and hypervariable regions.

Direct comparison with empirical human BCR datasets further confirmed the realism of the simulated repertoires. Amino-acid frequency distributions in simulated sequences closely matched those observed in real repertoires, with low Jensen–Shannon divergence, Hellinger distance, and total variation distance across all evolutionary depths examined. The modest decrease in divergence with increasing generation depth suggests that iterative rounds of mutation and selection progressively reinforce empirically observed compositional constraints, consistent with expectations from affinity maturation dynamics.

Importantly, simulated sequences also fully satisfied IMGT/JunctionAnalysis mutation acceptance criteria in junction regions. This compliance demonstrates that the implemented V(D)J recombination, junction modeling, and mutation processes preserve key structural and annotation-level constraints required for compatibility with standard immunogenomic analysis pipelines.

Beyond sequence-level realism, Ancestra produces lineage trees that capture essential qualitative features of B-cell evolution in germinal centers. Simulated lineages exhibited heterogeneous branching, non-uniform clonal expansion, and coexistence of multiple competing branches over extended evolutionary windows. Increasing the maximum generation depth led to deeper trees with greater terminal diversity, while preserving substantial internal branching rather than collapsing into linear trajectories. These properties are consistent with experimental observations that affinity maturation proceeds through complex, branching evolutionary processes rather than simple optimization paths.

A notable outcome of the lineage-explicit simulation is the emergence of multi-label lineage trees, in which identical nucleotide sequence states arise independently along distinct evolutionary paths. These events were observed in a substantial fraction of simulated lineages and occurred over both short and long topological distances within the tree. Importantly, these multi-path assignments do not reflect ambiguity in the simulated evolutionary process, but rather capture convergent outcomes arising from distinct mutational trajectories, highlighting the context-dependent nature of somatic hypermutation and selective pressures during antigen binding. From a methodological perspective, these structures provide a valuable testbed for evaluating how inference methods handle convergence, homoplasy, and non-unique mappings between sequences and evolutionary paths, phenomena that are difficult to assess using experimental data alone.

By integrating biologically grounded mutation and selection models with explicit lineage recording, Ancestra offers a flexible platform for benchmarking a wide range of repertoire-analysis tools. In particular, it enables systematic evaluation of lineage reconstruction algorithms, phylogenetic methods, and machine-learning approaches that incorporate evolutionary or lineage-level information. The availability of known ground truth allows performance to be assessed independently of algorithmic assumptions used during inference.

Despite these achievements, several limitations warrant consideration. In Ancestra, the sequences and lineage trees presented as output represent only a small fraction of the total diversity generated internally. During each generation, every surviving B cell produces multiple mutated candidates, most of which are evaluated and discarded because they fail selection criteria. This process can involve millions or even billions of candidate sequences over the course of a single simulation, which amplifies the computational cost of every design decision in the affinity evaluation.

To maintain scalability, affinity is computed using a fast, CDR3-centric alignment score rather than a full-sequence, biophysically detailed binding model. While this approach does not provide quantitative binding estimates, it captures the functional diversity necessary to drive selection and competition among clones, enabling biologically plausible evolutionary trajectories. The CDR3 captures the majority of functional diversity relevant for distinguishing competing clones during affinity maturation, and for our purposes we do not need an exact physical estimate of binding strength. The alignment score therefore serves as a computational proxy to generate consistent relative signals for selection and branching.

Importantly, these approximations are not hard-wired: users can choose whether to compute affinity using only the CDR3 or the full BCR sequence, and the affinity module is fully replaceable. Nevertheless, the current implementation focuses on heavy-chain evolution, does not explicitly model light-chain pairing, isotype switching, or spatial dynamics within germinal centers, and simplifies mutation and selection modeling relative to full biophysical realism. These trade-offs were imposed to allow lineage-explicit simulation of large, highly branching trees in a computationally tractable manner.

In future versions, we aim both to remove current limitations and to incorporate additional biological assumptions to increase realism and predictive accuracy. Planned enhancements include modeling BCR heavy chain sequences with multiple D segments, incorporating light-chain evolution, implementing more sophisticated binding and selection models, and enabling simulation of multi-founder germinal center reactions and interactions between parallel lineages. Furthermore, we plan to adapt the simulator for use across multiple species, enabling comparative studies of repertoire dynamics by incorporating non-human germline libraries and species-specific recombination parameters. These developments will allow Ancestra to maintain its lineage-explicit design while offering more biologically detailed simulations, enabling deeper investigation of immune repertoire evolution and more accurate benchmarking of lineage-aware analytical methods.

## 5. Conclusion

Ancestra provides a practical framework for generating AIRR-compatible synthetic BCR datasets in which the evolutionary record is preserved by design. By producing complete parent–child relationships, mutation traces, and standardized outputs alongside repertoire sequences, the simulator enables benchmarking tasks that are not feasible when lineage histories must be inferred indirectly from sequence collections.

A key outcome of this design is the ability to evaluate lineage-aware methods under controlled conditions, including cases where identical sequence states arise through distinct mutational paths. By making such events explicit in the ground truth, Ancestra supports more stringent and transparent testing of clonal grouping, phylogenetic reconstruction, and downstream evolutionary analyses.

Overall, Ancestra establishes a reproducible foundation for simulation-driven evaluation in immunogenomics and creates a clear path for future extensions that incorporate additional biological detail while retaining traceable lineage histories.

## 6. Data and Code Availability

All data and source code of Ancestra are publicly available at https://github.com/Reza-HZ/Ancestra.

