## Supplemental Figures for "Ancestra: A lineage-explicit simulator for benchmarking B-cell receptor repertoire and lineage inference methods"

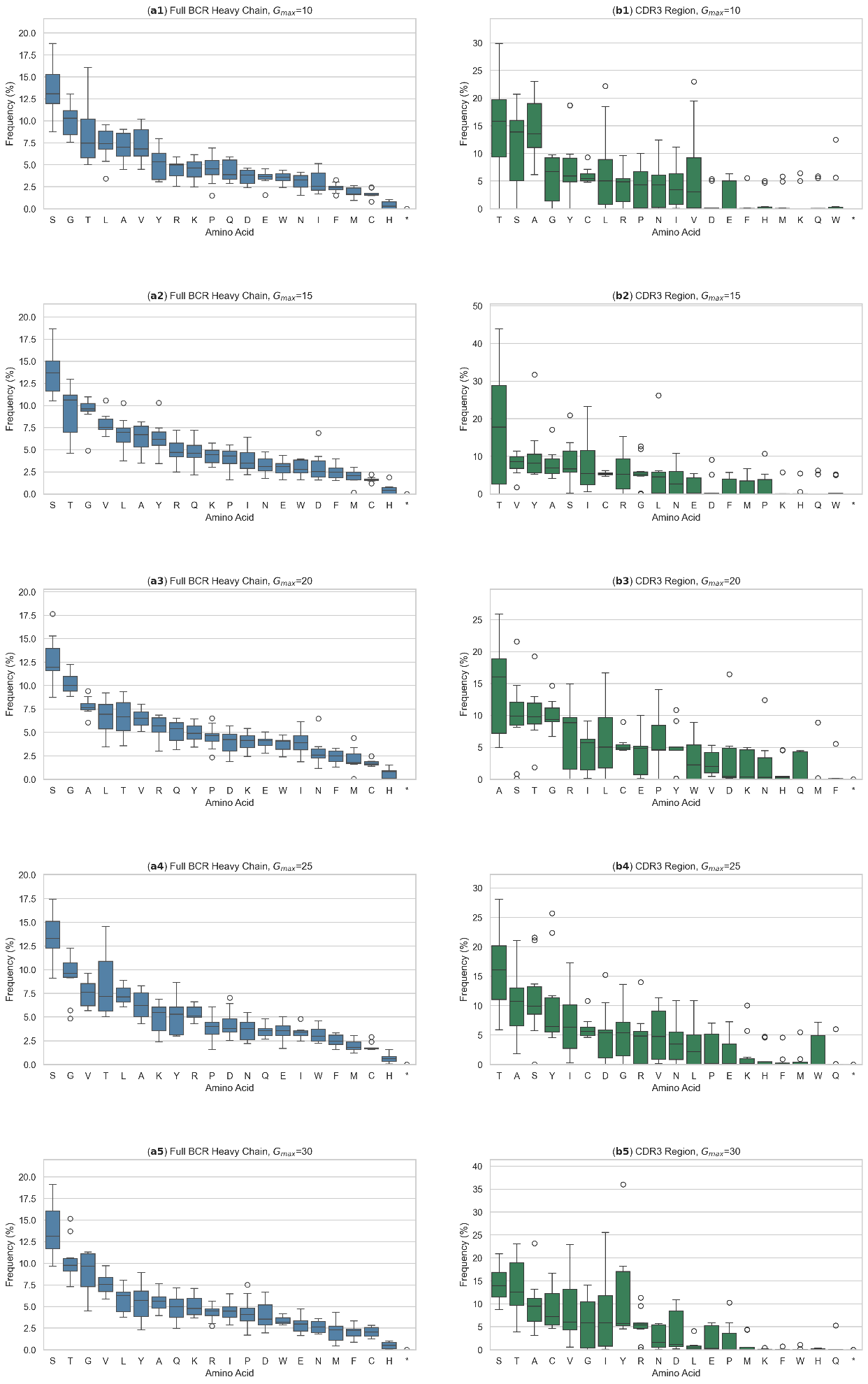

**Supplementary Figure 1.** Distribution of amino‑acid frequencies across simulated BCR heavy‑chain sequences, shown separately for each maximum generation depth ($G_{max}=\{10, 15, 20, 25, 30\}$). **(a1-a5)** Full‑sequence amino‑acid distributions remain stable across generations. **(b1-b5)** CDR3‑restricted distributions show a distinct compositional profile and greater heterogeneity than full sequences, consistent with region‑specific compositional biases encoded by the simulation framework.

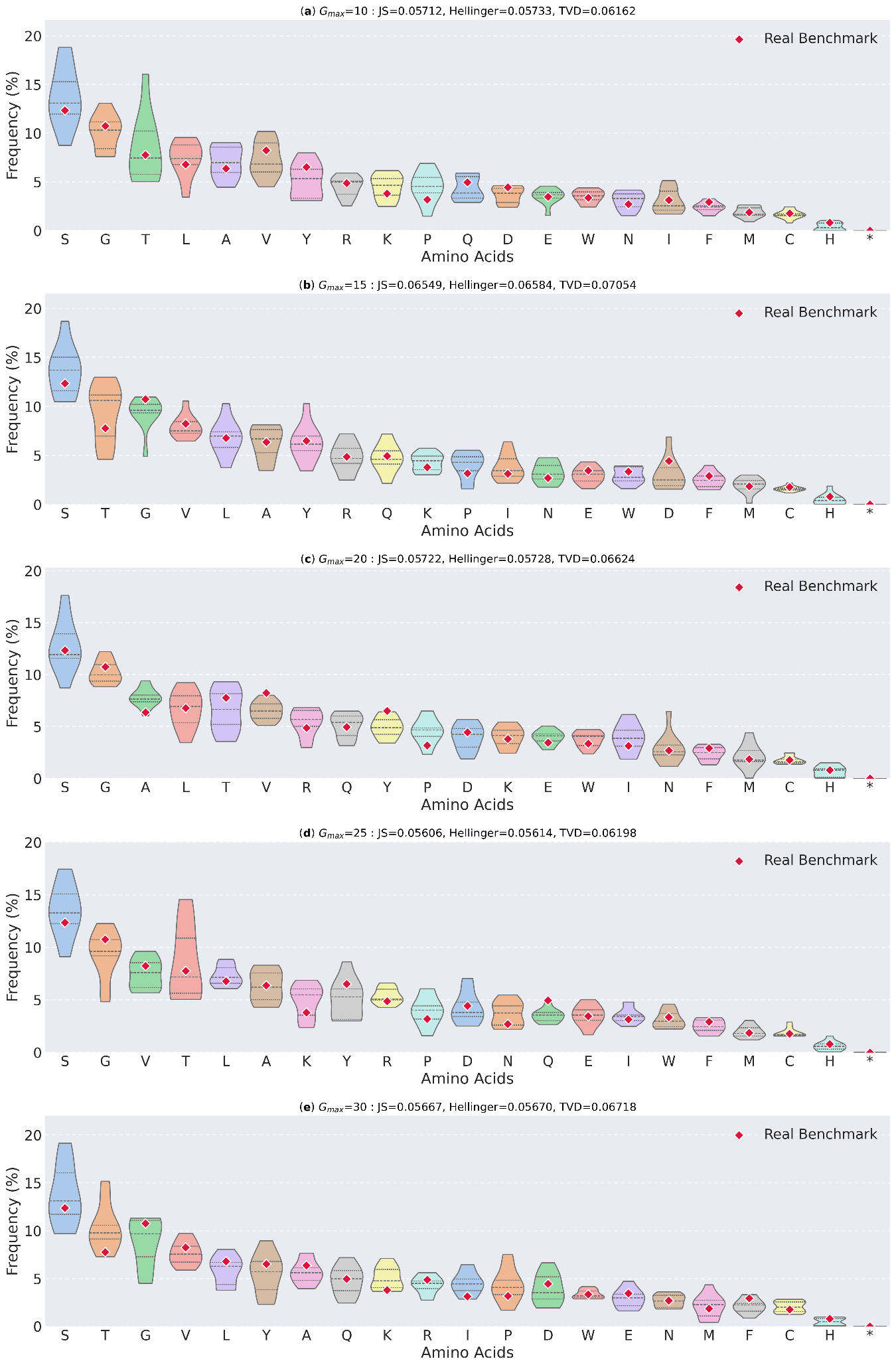

**Supplementary Figure 2.** Comparison of amino‑acid frequency distributions between simulated and real human BCR repertoires, shown separately for each maximum generation depth ($G_{max}=\{10, 15, 20, 25, 30\}$). Each violin represents across‑run variability of a single amino acid in the simulated repertoires; red diamond markers indicate corresponding frequencies in the real benchmark dataset (ABSD). For each generation depth, divergence metrics (Jensen‑Shannon divergence, Hellinger distance, and Total Variation Distance) are computed between the median simulated distribution and the empirical benchmark.
